# Repeatability of soma and neurite metrics in cortical and subcortical grey matter

**DOI:** 10.1101/2020.10.08.331595

**Authors:** Sila Genc, Maxime Chamberland, Kristin Koller, Chantal M.W. Tax, Hui Zhang, Marco Palombo, Derek K. Jones

**Author notes:** authors equally contributed to this work (Marco Palombo, Derek K. Jones).

## Abstract

Diffusion magnetic resonance imaging is a technique which has long been used to study white matter microstructure *in vivo*. Recent advancements in hardware and modelling techniques have opened up interest in disentangling tissue compartments in the grey matter. In this study, we evaluate the repeatability of soma and neurite density imaging in a sample of six healthy adults scanned five times on an ultra-strong gradient magnetic resonance scanner (300 mT/m). Repeatability was expressed as an intraclass correlation coefficient (ICC). Our findings reveal that measures of soma density (mean ICC=.976), neurite density (mean ICC=.959) and apparent soma size (mean ICC=.923) are highly reliable across multiple cortical and subcortical networks. Overall, we demonstrate the promise of moving advanced grey matter microstructural imaging towards applications of development, ageing, and disease.

## 1 Introduction

Conventional T1-weighted magnetic resonance imaging (MRI) is a useful tool in determining clinically relevant regional differences in grey matter volume, cortical thickness, surface area and gyrification. However, these crude macroscopic measures do not provide information on which distinct cellular features (e.g. cell bodies and neurites) and packing configurations drive differences in macroscopic measures. Diffusion MRI (dMRI) can enhance sensitivity to much smaller structures by probing water diffusion that is modulated by the presence of micrometer-scale compartments. Previous studies have applied the commonly used diffusion tensor imaging (DTI) technique to profile microstructure in the grey matter [e.g. 1, 2], however biological interpretations are limited as DTI metrics are non-specific to the aforementioned microstructural compartments.

Progress in acquisition and modelling methods using ultra-strong gradient and ultra-high b-value dMRI [3, 4] hold promise for disentangling and quantifying biologically meaningful cellular components *in vivo* [5, 6]. One recent model-based method to study grey matter microstructure is Soma and Neurite Density Imaging (SANDI)[5], which aims to disentangle microstructural contributions from cellular projections (neurites: including axons, dendrites and glial processes), soma (neuronal cell bodies and glia) density and their apparent size, and extra-cellular space.

The original SANDI paper demonstrated results in humans using ultra-high b-value data (up to 10,000 s/mm^2^) [5]. In this study, we utilise a rich repeatability database of scan-re-scan dMRI data acquired from 6 healthy participants, each across 5 sessions [7] on an ultra-strong gradient MR scanner [3, 4]. Our primary aim is to establish whether SANDI metrics are repeatable at lower b-values (up to 6,000 s/mm^2^) to establish the translatability and utility of advanced microstructural imaging in cortical and subcortical grey matter.

## 2 Methods

### 2.1 Image acquisition and pre-processing

The data used for this study were previously reported by Koller at al. [7], comprising a sample of 6 healthy adults (3 female) aged 24-30 years. This study was approved by a local ethics board. Each participant was scanned five times in the span of two weeks on a 3.0T Siemens Connectom system with ultra-strong (300 mT/m) gradients.

Structural data were acquired using a magnetization-prepared rapid acquisition with gradient echo (MPRAGE, voxel-size = 1*×* 1 *×*1 mm) and multi-shell dMRI data were collected (TE/TR = 59/3000 ms; voxel size = 2 *×*2 *×*2 mm; b-values= 0 (14 vols), 200;500 (20 dirs), 1200 (30 dirs), and 2400;4000;6000(60 dirs) s/mm^2^). dMRI data were acquired in an anterior-posterior (AP) phase-encoding direction, with additional b=0 s/mm^2^ images acquired in the PA direction. Pre-processing involved: noise estimation using Marchenko-Pastur Principles Component Analysis (MP-PCA) [8] and subsequent denoising in *MRtrix3* [9], correction for signal drift [10], motion, eddy, and susceptibility-induced distortions [11, 12], gradient non-linearities [13, 14], Gibbs ringing artefacts [15], and bias field [9, 16].

### 2.2 Image processing and analysis

The SANDI compartment model was fitted to the pre-processed dMRI dataset for each subject using the machine learning approach described in [5], based on random forest regression. Four parameters of interest were investigated:

i. the intraneurite signal fraction, *f*_*intraneurite*_
ii. the intrasoma signal fraction, *f*_*intrasoma*_
iii. the soma radius, *R*_*soma*_ (*µ*m)
iv. the extracellular signal fraction, *f*_*extracellular*_

Additionally, an estimate of model uncertainty was obtained using the quartile deviation of predictions (QD) from the ensemble of regression trees. To complement the SANDI estimates, diffusion tensor estimation was performed on the b=1200 s/mm^2^ shell using an iteratively reweighted linear least squares estimator. The tensor-derived parameters fractional anisotropy (FA) and mean diffusivity (MD) were computed.

T1 data were co-registered to an upsampled b=0 s/mm^2^ image (1mm isotropic) and processed through Freesurfer [17] to obtain cortical and subcortical parcellations using the Destrieux atlas [18]. This resulted in 74 cortical regions per hemisphere, alongside subcortical regions. We studied seven different functionally-defined networks from the Yeo functional network atlas [19] (Figure 1). The subcortical parcellation was treated as a single sub network, resulting in eight total subnetworks for each participant for further statistical analysis. Follow-up analyses of individual subcortical regions were restricted to the amygdala, caudate, hippocampus, pallidum and thalamus. Network labels (L) were resampled to each individual subject’s diffusion space, and we computed the intersection between the cortical ribbon (R) and resampled network labels (L*∩*R).

**Fig. 1.**
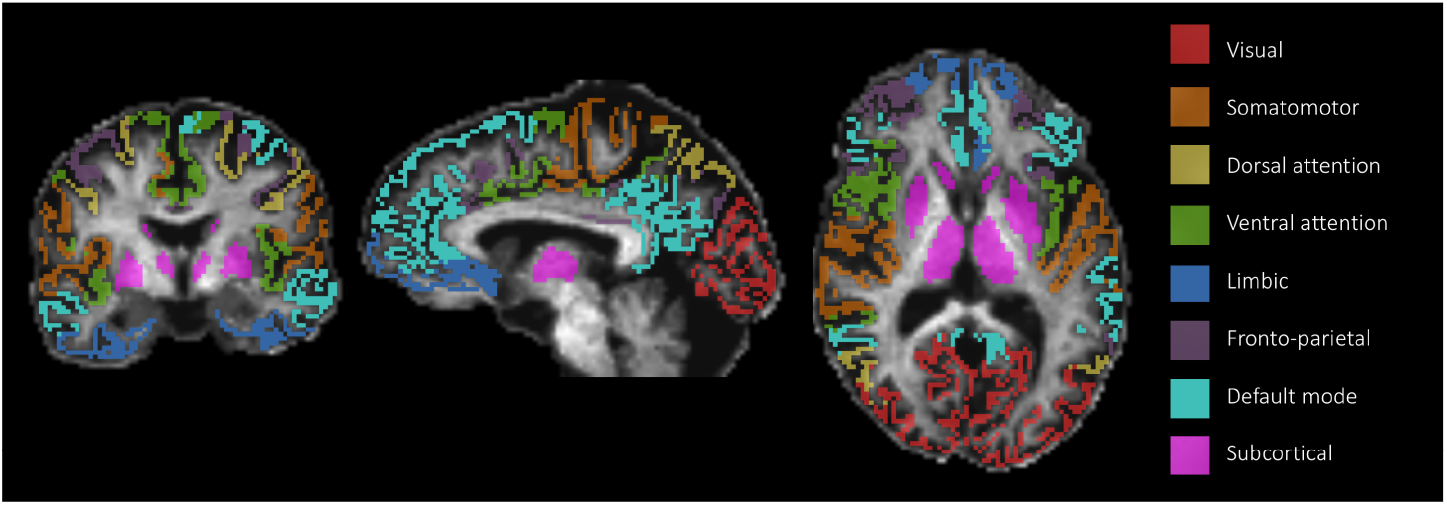
A representation of the eight cortical and subcortical sub-networks [19] on a representative participant

Statistical analyses were performed within R (v3.4.3) and RStudio (v1.2.1335). The intra-class correlation coefficient (ICC; two-way random effects, absolute agreement) was computed for assessment of test-re-test repeatability of SANDI and DTI metrics (Table 1). Summary statistics were computed using an analysis of variance (ANOVA), and lower and upper estimates of each ICC represent the bounds of the 95% confidence interval (CI). Based on the number of comparisons (8 networks x 6 metrics = 48 comparisons) we adjusted our p-value threshold of significance using a Bonferroni correction to *p <* .001.

**Table 1.**
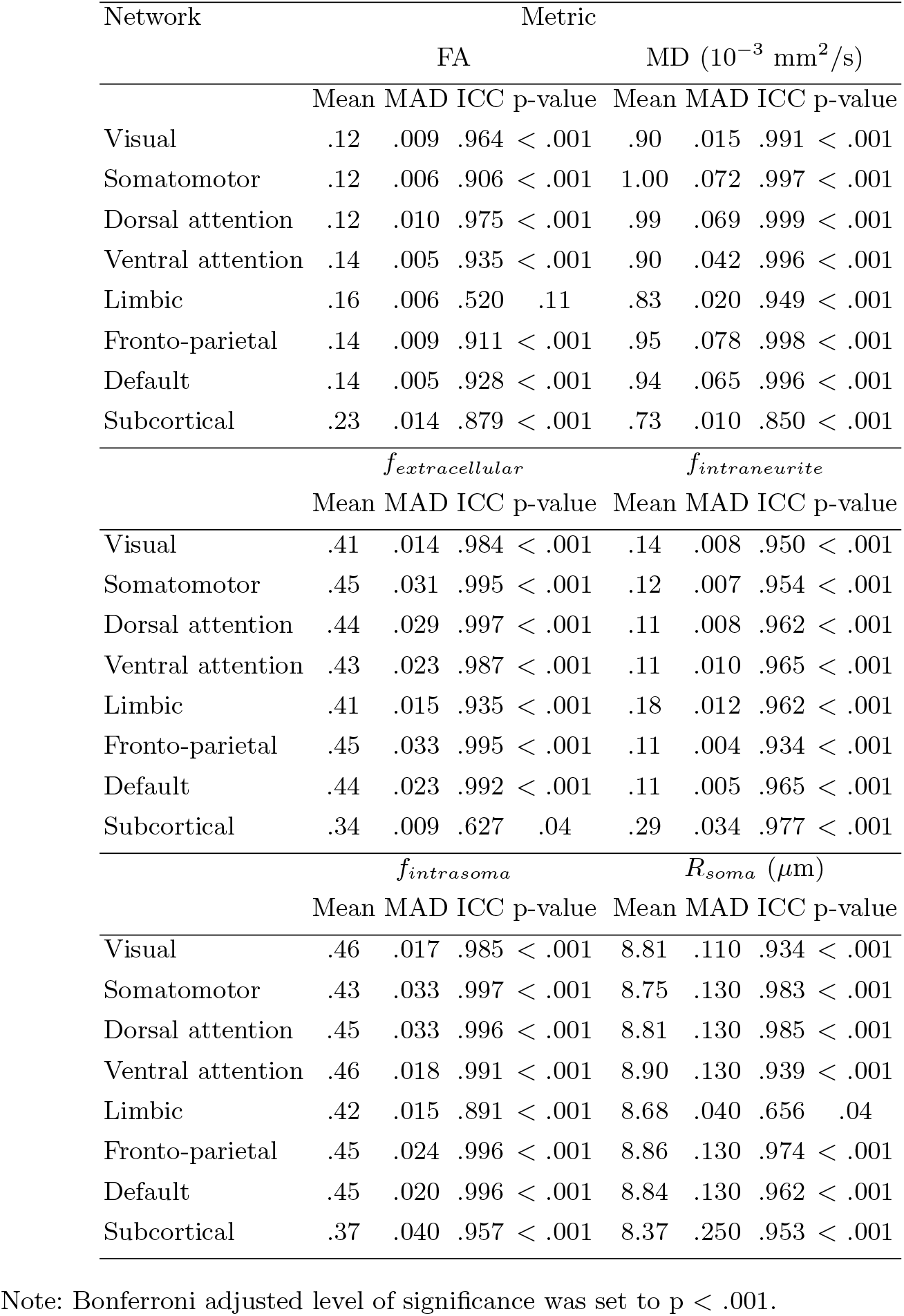
Statistics on test-re-test repeatability of DTI and SANDI metrics. Statistics summarise the mean, median absolute deviation (MAD), intra-class correlation coefficient (ICC) and p-value across all repeated measurements

## 3 Results

The results of the repeatability analysis and estimated values for *f*_*intraneurite*_, *f*_*intrasoma*_, *R*_*soma*_, and *f*_*extracellular*_ are reported in Table 1. These values were comparable to previously reported values estimated using ultra-high b-value data [5]. Intra-subject variability was generally very low for all metrics across all grey matter networks (Figures 2 & 3), reflected by high ICC values for SANDI metrics (mean ICC=.95) and DTI metrics (mean ICC=.93). Regions and metrics with lower repeatability and greater intra-subject variability included FA in the limbic network (ICC=.52, p=.11), *f*_*extracellular*_ in the subcortical grey matter (ICC=.63, p=.04) and *R*_*soma*_ in the limbic network (ICC=.66, p=.04).

**Fig. 2.**
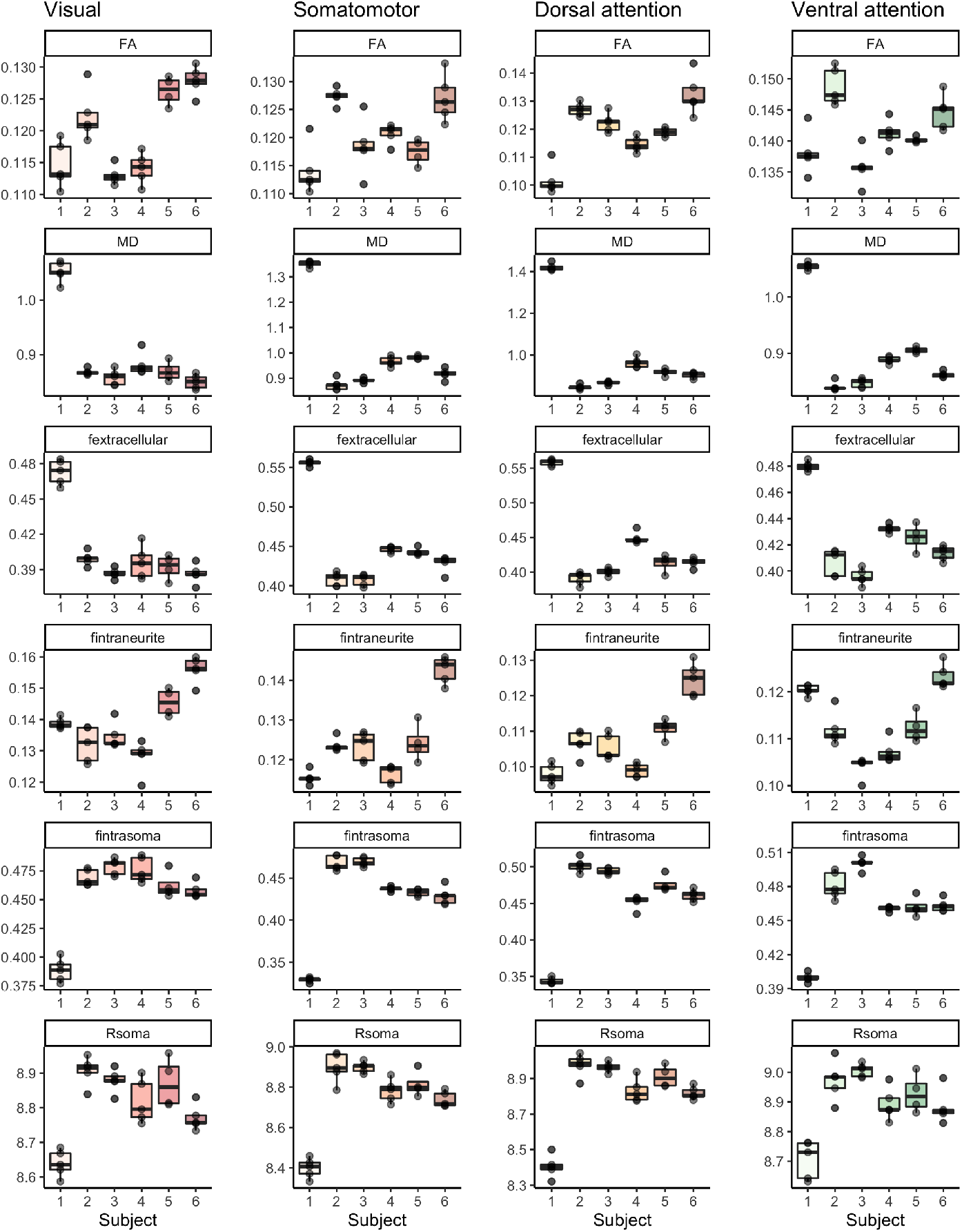
Spread of values for DTI and SANDI metrics in networks 1-4. Each subject (on the x-axis) has 5 data points representing each scan. The y-axis represents the point estimate of each microstructural metric: FA, MD (10^*−*3^ mm^2^/s), *f*_*extracellular*_, *f*_*intraneurite*_, *f*_*intrasoma*_, and *R*_*soma*_ (*µ*m)

**Fig. 3.**
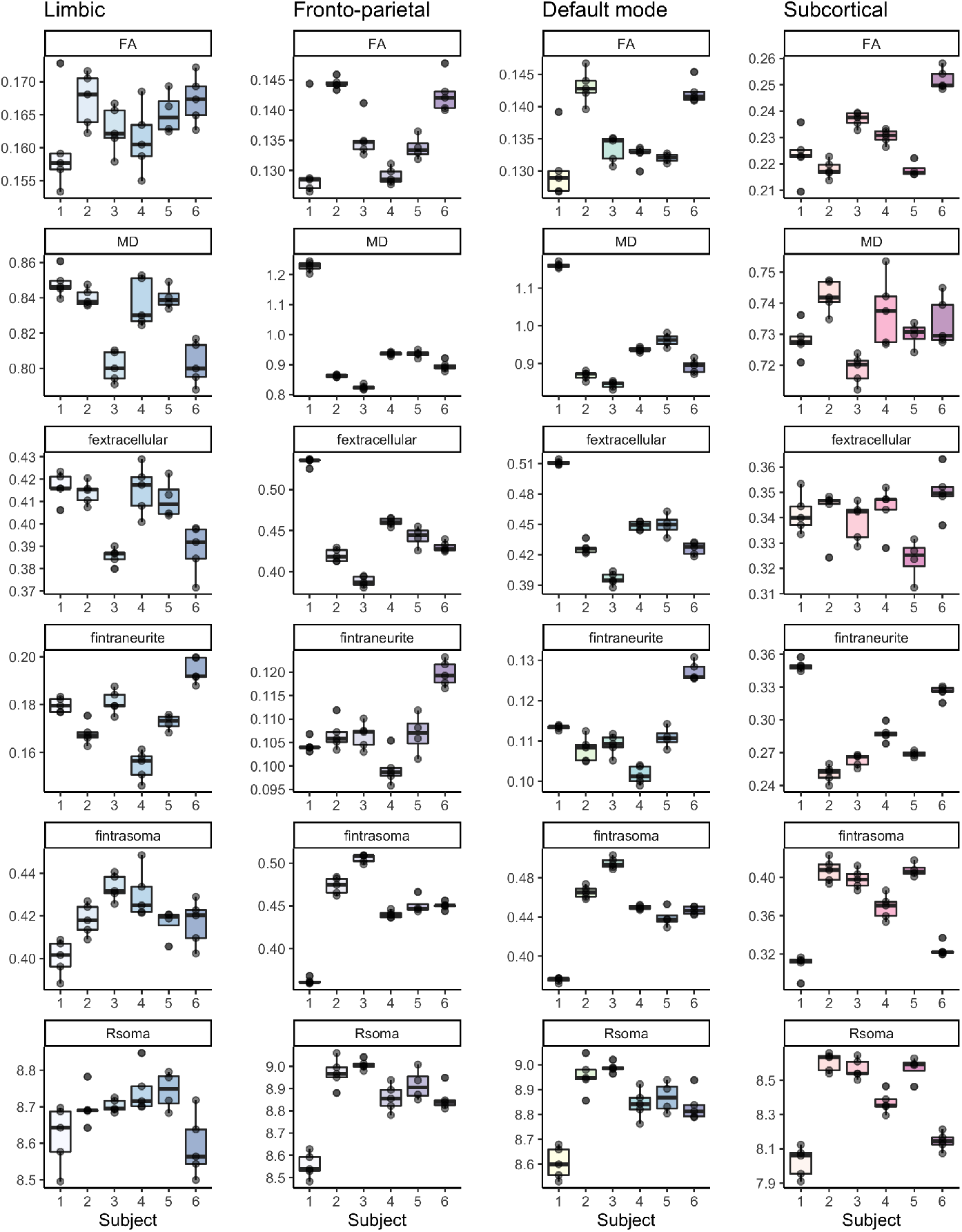
Spread of values for DTI and SANDI metrics in networks 5-8. Each subject (on the x-axis) has 5 data points representing each scan. The y-axis represents the point estimate of each microstructural metric: FA, MD (10^*−*3^ mm^2^/s), *f*_*extracellular*_, *f*_*intraneurite*_, *f*_*intrasoma*_, and *R*_*soma*_ (*µ*m)

Despite high repeatability across both DTI and SANDI metrics in the grey matter, DTI metrics exhibited larger uncertainty around ICC estimates, indicated by larger error bars (Figure 4a). In terms of SANDI model uncertainty, ICC values in the limbic network for all QD estimates had a wide variation indicated by larger error bars for the bounds of each ICC estimate (Figure 4b), and similar patterns were observed for *f*_*intrasoma*_ in the subcortical grey matter.

**Fig. 4.**
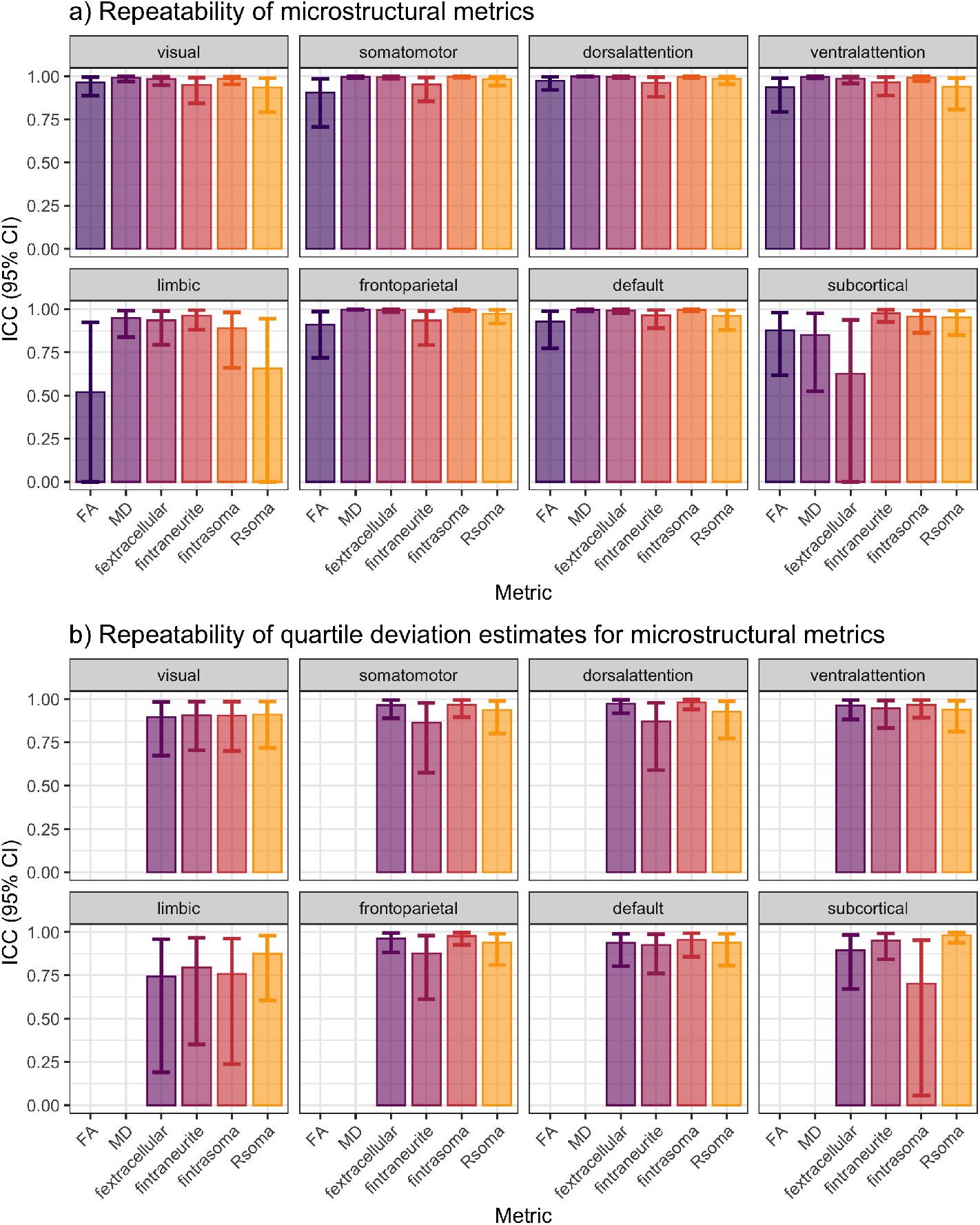
Intra-class correlation coefficients (two-way random effects, absolute agreement) for test-re-test repeatability of a) DTI and SANDI metrics, and b) the quartile deviation of SANDI metrics, in cortical and subcortical grey matter networks. Error bars represent the bounds of the 95% confidence interval (CI) for each ICC estimate

The results of the regional subcortical analysis are presented in Figure 5. We observed low repeatability of *f*_*extracellular*_ in all regions apart from the left amygdala and left caudate (Figure 5a,b). For *R*_*soma*_, only the left amygdala, left pallidum, and right hippocampus exhibited low repeatability (Figure 5a,b). The distribution of QD estimates for *f*_*intrasoma*_ suggest potential variation in model fit between regions (Figure 5c).

**Fig. 5.**
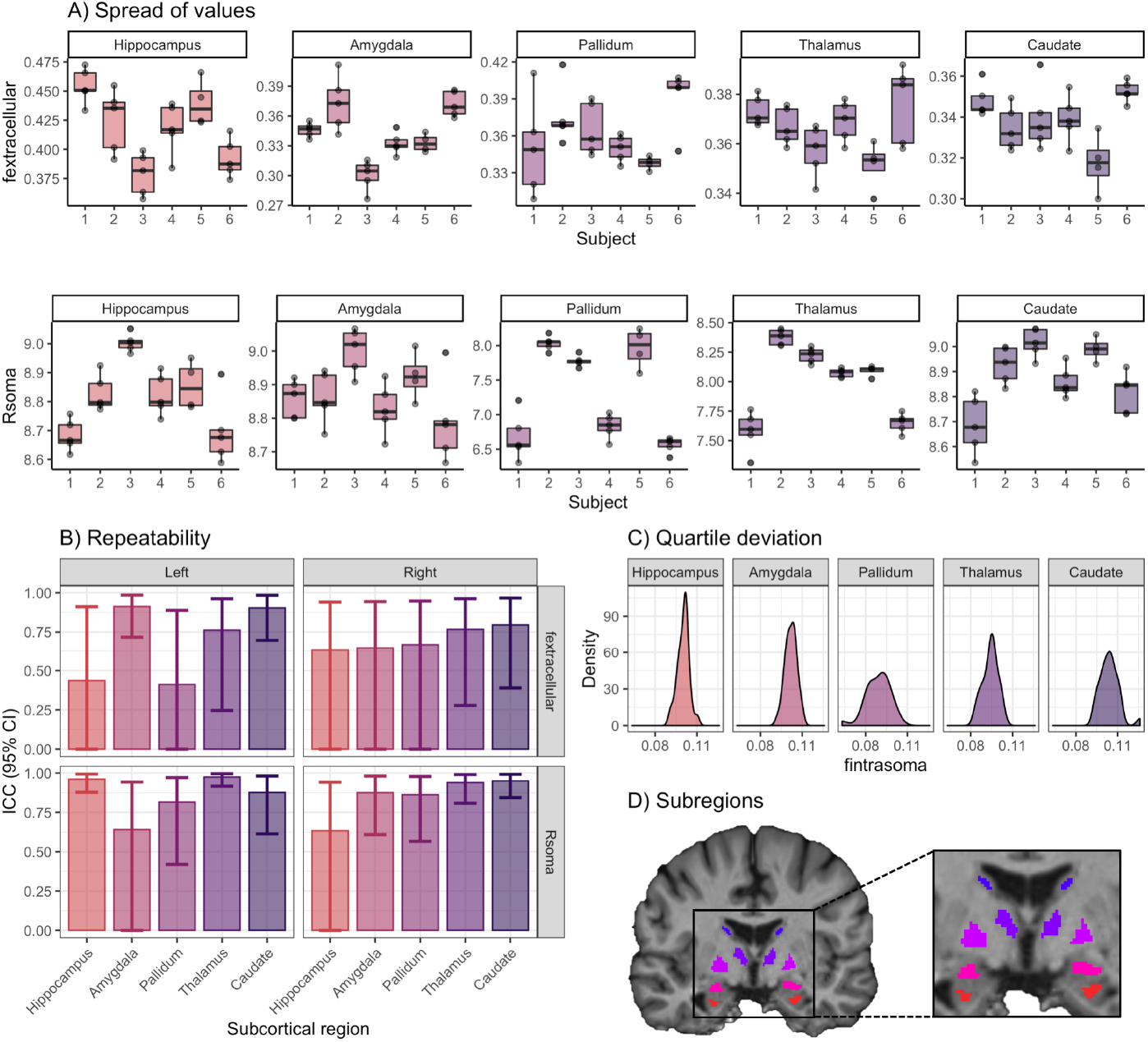
Regional analysis in subcortical grey matter for the hippocampus, amygdala, pallidum, thalamus and caudate. A) Spread of *f*_*extracellular*_ and *R*_*soma*_ values in the left hemisphere; B) Intra-class correlation coefficients (two-way random effects, absolute agreement) for test-re-test repeatability of *f*_*extracellular*_ and *R*_*soma*_; C) Distribution of quartile deviation (QD) estimates for *f*_*intrasoma*_; D) Regions of interest used in the analysis, obtained using Freesurfer [17] and eroded by 1 mm

## 4 Discussion

Our findings reveal that estimates of grey matter microstructure using soma and neurite density imaging are highly stable across repeated imaging sessions. We demonstrate high repeatability in a number of functional networks, known to share structural covariance [19]. In addition, the soma signal fraction variation across limbic and visual networks follows the estimated anterior to posterior gradient of cell density in the cortex of human and other primates [20]. Overall, our findings of high repeatability of dMRI metrics in the grey matter suggest the increased power to detect group differences in applications of this technique.

The limbic network showed consistently lower repeatability and model uncertainty amongst both DTI and SANDI metrics. Given the anatomical location of these fronto-temporal structures, it is likely that susceptibility-induced distortions may detrimentally influence the repeatability of certain diffusion MRI metrics. The effect of gradient non-linearities and spatiotemporally varying b-values could impact repeatability, if the subject is placed in a slightly different position in the scanner. Despite this general observation of the SANDI metrics studied here, only soma radius exhibited low repeatability in this network.

Therefore, *R*_*soma*_ estimates in fronto-temporal structures should be interpreted with caution, particularly in populations where these artefacts may be exaggerated (e.g. fronto-temporal dementia).

Upon further analysis of individual subcortical regions, the repeatability of *f*_*extracellular*_ was generally low. Tissue properties within subcortical regions are heterogeneous, as sub-segments can differ in their neurite and soma composition [21, 22]. These anatomical variations may influence the estimates reported here, and as such, even finer parcellation of individual subcomponents would be an important avenue of future research.

Finally, we have demonstrated that SANDI estimates obtained from moderate-to-high b-values (up to b=6000 s/mm^2^) are comparable in terms of magnitude to previous estimates derived from ultra-high b-values (e.g. up to b=10,000 s/mm^2^). Whilst a direct comparison between multiple sampling schemes tested on the same participant across repeated scans would be required to confirm similar magnitudes of repeatability, based on our findings we are confident that the repeatability is high enough to be acceptable for research applications using high b-values. Now that we have established that these novel markers of grey matter microstructure are stable across repeated sessions, the next step is to pinpoint the underlying tissue properties driving rapidly changing grey matter macrostructure, such as that observed in neurodevelopment and neurodegeneration.

